# Language models annotate mass cytometry clusters within the range where human experts disagree

**DOI:** 10.64898/2026.10.20.760379

**Authors:** Oliver Hunewald, Erica T. Grant, Theresa-Maria Boehm, Annette Kuehn, Vladimir Despotovic, Mahesh S. Desai, Petr V. Nazarov

**Affiliations:** Bioinformatics & AI, Department of Medical Informatics, Luxembourg Institute of Health, 1A-B rue Thomas Edison, L-1445 Strassen, Luxembourg; Department of Infection and Immunity, Luxembourg Institute of Health, 29 rue Henri Koch, L-4354 Esch-sur-Alzette, Luxembourg; Department of Cancer Research, Luxembourg Institute of Health, 1A-B rue Thomas Edison, L-1445 Strassen, Luxembourg; Faculty of Science, Technology and Medicine, University of Luxembourg, L-4354 Esch-sur-Alzette, Luxembourg

## Abstract

Cell-type identification of clusters from unsupervised cytometry pipelines (e.g. FlowSOM, GigaSOM) is a time-consuming, expert-dependent bottleneck, typically requiring an expert to read a heatmap of median marker intensities and assign a label to each cluster. Large language models (LLMs) can annotate single-cell RNA-seq clusters from marker-gene lists, but cytometry cluster data differ in both dimensionality and data type. We present a direct procedure in which cluster medians are scaled per marker and grouped by similarity into a few dozen metaclusters. The continuous marker profiles are then passed to the LLM, which returns a cell-type label and confidence level for each metacluster in a single call, without a gating hierarchy, manually defined marker-specific thresholds, or task-specific training data. Across four human and mouse datasets spanning blood, bone marrow and gut, a single well-chosen model reproduces the reference annotation at the major-lineage level with 85–98% metacluster-level concordance (cloud Claude Sonnet 4.6 85–98%, best local model 83–96%), substantially exceeding majority-class baselines; chance-corrected agreement is Cohen’s *κ* = 0.67– 0.83. Local open-weight models perform competitively with the cloud model on the independent benchmarks, enabling application to clinical or patient-derived data subject to data-protection restrictions. Reliability is model-dependent: no local model is best everywhere, and the frontier model Claude Opus 5 was no more reliable than Claude Sonnet 4.6, the most consistent in concordance across datasets. Because no annotation is absolute ground truth, we obtained two additional expert annotations of the same continuous heatmaps, blinded to the reference and to all model output, to establish a method-matched agreement ceiling. The experts agreed on *∼*99% of major-lineage assignments but only *∼*73% of T-cell maturation states; on that axis the best models fall within the expert disagreement range. LLM-based annotation is therefore a practical, training-free alternative to rule-based annotation, provided its output is checked against the intended panel and species.

## 1 Introduction

High-dimensional cytometry by time-of-flight (CyTOF, or mass cytometry) measures 30–50 markers per cell across millions of cells. In addition to classical manual gating, in which cell populations are sequentially identified using gates on biaxial plots, modern cytometry workflows increasingly use unsupervised clustering to identify cellular populations, with self-organising-map approaches such as FlowSOM being widely adopted [1]. Cell-type identification of the clusters requires an expert to inspect each cluster’s expression profile, typically presented as a heatmap of median marker expression. This step is slow, subjective, hard to reproduce, and it is the practical bottleneck in most cytometry workflows [2, 3]. We evaluate whether a large language model (LLM) can substitute for that expert step. However, cell-type annotation is inherently interpretive, and no universally accepted ground truth exists against which model performance can be evaluated.

Two lines of work aim to automate annotation. Supervised and deep-learning annotators (e.g. Scyan [4], ImmCellTyper [5], and human-level automated gating [6]) require training data, labelled examples, or an explicitly encoded panel. Separately, LLMs have been applied to single-cell RNA-seq annotation from marker-gene lists (GPTCelltype [7]; the LLM mode of CytoAnalyst [8]). Marker-matching methods such as Scyan and ImmCellTyper can annotate cell populations directly from continuous protein-expression profiles without explicit gating. However, these approaches rely on a pre-specified, panel-specific reference that encodes expert knowledge of marker–cell-type associations and therefore requires prior definition for the panel under analysis. Ontology-based labelling assigns standardised names to populations from their marker signatures [9, 3]. It depends on a curated ontology and can return only the terms that ontology defines. The LLM’s role here is to supply the marker-to-cell-type knowledge on demand, per dataset, rather than requiring it to be encoded by hand. Marker-to-cell-type correspondence is densely documented in the immunology literature, and LLMs trained on that literature encode substantial biomedical knowledge [10]. The scRNA-seq results demonstrate that this is already sufficient to reproduce manual annotation from marker-gene lists. What is untested is whether it transfers to continuous protein intensities, and whether labels are driven by marker identity rather than by the numerical pattern of expression values alone.

Cytometric panels are designed a priori to capture markers informative of cellular lineage and functional state, which experts then interpret based on their continuous intensities. The LLM receives each metacluster’s continuous per-marker profile and returns a cell-type label and confidence, with no task-specific training data and no manually encoded panel-specific reference. The procedure is deliberately minimal, emphasising evaluation of the annotator over model or prompt engineering. Our contributions are as follows:

1. An evaluation framework for annotators without a ground truth. Two experts independently annotate the same continuous profiles the model receives, giving a method-matched inter-expert agreement ceiling (§3.3). Inter-annotator agreement as a ceiling is established practice in natural language processing [11] and medical-image reader studies [12]; here, we extend this approach to cytometry annotation.
2. A comprehensive evaluation across four datasets, seven models, and three annotation paradigms, spanning human and mouse data from blood, bone marrow and gut, and including two public benchmarks independent of our laboratory, tools, and annotators (§3.2).
3. A minimal, training-free annotation procedure mapping continuous metacluster profiles to hierarchical cell-type labels with a confidence level in a single LLM call (§2.4). We further characterize the reliability of the approach and demonstrate its feasibility with local open-weight models for privacy-restricted data (§3.4).

## 2 Materials and methods

### 2.1 Datasets

We used four datasets from published studies, spanning three distinct annotation paradigms (Table 1). One was annotated with **CyCadas** [13], a hierarchical, threshold-based annotation algorithm that assigns each cluster a cell type by walking it through a tree of per-marker binary (positive/negative) threshold calls. This is **Capelle2023**, a Parkinson’s-disease whole-blood study [14] with 27 markers, for which the 1,600-cluster solution and CyCadas annotations were obtained from [13].

**Table 1:** Summary of the four datasets, spanning three annotation paradigms (CyCadas, independent manual gating, and direct expert heatmap inspection), two species, and three tissues (blood, bone marrow and gut). “Ref. method” indicates how the reference annotation was produced. Capelle2023’s 31 reference populations resolve lineage plus T-cell coreceptor and maturation subsets. Parrish2023’s “58 / 26” denotes 58 fine populations that the original authors grouped into 26 coarse clusters.

| Dataset | Species | Tissue | Clusters | Markers | Ref. populations | Ref. method |
| --- | --- | --- | --- | --- | --- | --- |
| Capelle2023 | Human | Whole blood | 1,600 | 27 | 31 | CyCadas |
| Levine2015 | Human | Bone marrow | 100 | 32 | 14 | Manual gating |
| Samusik2016 | Mouse | Bone marrow | 100 | 39 | 24 | Manual gating |
| Parrish2023 | Mouse | Colonic lamina propria | 400 | 27 | 58 / 26 | Direct heatmap |

Two are independent public benchmarks obtained through the **HDCytoData** Bioconductor package [15] (originally deposited in FlowRepository under FR-FCM-ZZPH), both with manual-gating reference annotations. They were chosen precisely because they are independent of the CyCadas dataset in annotation tool, laboratory, and annotator. **Levine2015** [16] is human bone marrow with 32 markers and 14 manually gated populations; **Samusik2016** [17] is mouse bone marrow with 39 markers and 24 manually gated populations. For these two datasets, no pre-existing cluster table was available. The upstream fine-clustering step was therefore performed here to match the procedure applied to the other two datasets: labelled single-cell data were arcsinh-transformed and partitioned into 100 clusters using *k*-means. Because this step only has to yield lineage-pure fine clusters for the LLM to label, and fine-cluster purity against the reference labels was 96–100%, the choice of fine-clustering algorithm does not cap achievable concordance.

Finally, **Parrish2023** is a published CyTOF dataset of mouse colonic lamina propria-derived cells [18], comprising 400 FlowSOM clusters, 27 markers, and 58 fine populations that the original authors further grouped into 26 coarse UMAP-level clusters while retaining functionally distinct subsets, such as Th1/Th2/Th17/Treg and ILC1/2/3. Its reference annotations were generated by direct expert inspection of marker-expression heatmaps. We annotate the study’s specific-pathogen-free (SPF) subset (Parrish et al., Extended Data Fig. 5), which is deposited on Zenodo [19]. This dataset therefore represents a third annotation paradigm, distinct from both CyCadas-based threshold annotation and manual gating, and most closely matches the LLM annotation task itself. It also extends the evaluation to tissue-resident populations, including innate lymphoid cells and T-helper subsets, that are not represented in peripheral blood. Although some authors of the present study contributed to Parrish2023, its annotation is independent of CyCadas in tool, annotator, and paradigm. Cross-lab independence, on the other hand, comes from Levine2015 and Samusik2016, which share no institution, tool, or annotator with this work.

Sample counts, cohort characteristics and per-sample cell counts for all four datasets are given in Table S1.

### 2.2 Per-marker scaling

Even after arcsinh transformation, each marker spans a different dynamic range, so absolute “high” and “low” values are not comparable across markers without normalisation. Scaling each marker to a common [0, 1] range ensures a given value carries a consistent interpretation across markers, and reproduces what an expert sees on a heatmap, where each marker’s colour is scaled to its own range. Each marker is clipped at its 1st and 99th percentiles across clusters and then min-max scaled to [0, 1]. Percentile-rank scaling is avoided because it maps each marker onto a uniform distribution, expanding background variation in low-signal markers to fill the full range. Clipping preserves the relative spacing of intensities, so weakly expressed markers remain low after scaling.

### 2.3 Metaclustering

Clusters were grouped into a fixed number of metaclusters using deterministic Ward hierarchical clustering on the scaled profiles, with each metacluster represented by the frequency-weighted mean profile of its constituent clusters. Concordance results are scored at the major-lineage level (§2.6), except for the fine-attribute ceiling analysis (§3.3).

The number of metaclusters therefore only needs to be sufficient to avoid merging distinct lineages. For example, the 40 metaclusters used for Parrish2023 readily resolve its seven major lineages and 26 coarse groups without requiring separation of all 58 fine populations. The count is set to approximate the granularity of the available reference annotation, using 40 metaclusters for all four datasets, as a conservative, untuned choice. Lineage-level purity relative to the reference is ∼99% across metacluster counts, indicating that metaclusters are not substantially lineage-mixed.

### 2.4 Direct LLM annotation

The proposed LLM-based cytometry annotation workflow is summarised in Fig. 1. Each metacluster profile is serialised as a compact list of bright (≥ 0.6), intermediate (0.35–0.59), and low markers, together with the population size. The LLM is prompted to interpret the profile as an expert immunologist reading a continuous heatmap, with the tissue and species specified and intermediate expression explicitly treated as biologically informative (e.g. CD56-dim NK cells). For each metacluster, it returns a JSON record with a cell-type label and a confidence level. With 40 metaclusters per dataset, below the batch limit of 50 (see Models), all metaclusters of each dataset are annotated in a single call. Labels are then propagated to the constituent clusters, and the sampling temperature is set to 0. The method requires no gating tree, no hand-authored per-marker thresholds, and no training data.

**Figure 1:**
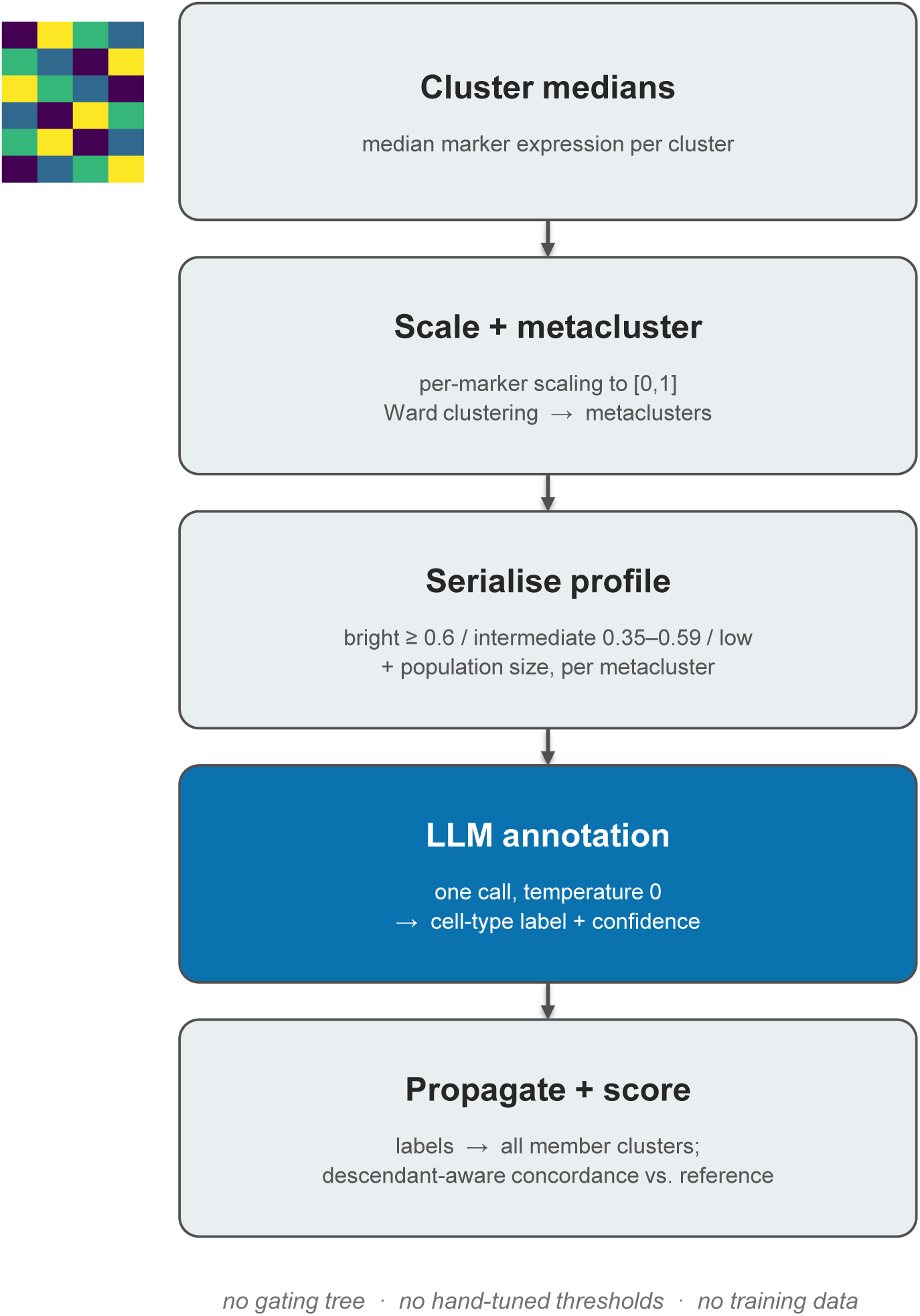
The direct-annotation pipeline. Cluster medians are scaled per marker, grouped into a few dozen metaclusters, and each metacluster’s continuous profile is serialised as a bright/intermediate/low marker list and sent to the LLM in a single call; the returned labels propagate to the constituent clusters and are scored descendant-aware against the reference.

### 2.5 Models

We evaluated one cloud model, Claude Sonnet 4.6 (claude-sonnet-4-6), and six local open-weight models available as of July 2026, limited to approximately 128B parameters to accommodate the available hardware: gpt-oss 120B, qwen3.6 35B (MoE), gemma4 31B, gemma3 27B, Llama 4 Scout, and medgemma 27B, a medically fine-tuned gemma3 model. All local models were served through Ollama using an OpenAI-compatible endpoint; model versions, tags, precision and on-disk sizes are given in Table S2.

Only the qwen3.6 tag pins a quantisation (bf16); the remaining local models use default distribution tags quantised to approximately 4 bits per parameter, so the local models were not served at matched numerical precision. For all local models the client specified a temperature of 0 and no other generation parameters, leaving remaining sampling and context settings at the server defaults. Metaclusters were submitted in batches of up to 50 per call, a client-side, hardware-dependent setting chosen to avoid time-outs and not imposed by the models or the method. Prompts were under 6k tokens for every dataset, well within the 131,072-token context limit allocated per model (128,000 for MedGemma, as specified by its distribution tag). Server logs confirmed that no prompt truncation occurred. Local models were served on a single NVIDIA GB10 (Grace Blackwell) node with 119 GiB of unified memory, with model weights fully resident on the GPU.

As a **frontier-model control**, Claude Opus 5 was additionally evaluated on all four datasets to determine whether a more capable cloud model provides improved annotation performance. Opus 5 was run at its default sampling because the API rejects the temperature parameter for this model; all other models used temperature 0. Three runs were performed per dataset; the mean is reported, together with the run-to-run range and lineage-level stability (§3.5). Note that temperature 0 makes decoding greedy but does not guarantee identical output: batching, hardware and serving-stack non-determinism can still change which token is selected at near-ties, so run-to-run agreement below 100% is expected.

### 2.6 Evaluation

LLM annotations are evaluated by comparison with existing expert annotations of the same data. None of the available references, whether produced by a threshold-based tool, a manual gating hierarchy, or direct expert interpretation of a heatmap, is treated as a gold standard: each represents a human judgement obtained through a particular annotation procedure, and they can disagree. Evaluation is therefore framed as concordance between annotations rather than accuracy against a fixed reference, and in §3.3 model concordance is calibrated against the agreement between two independent human experts. For scoring, labels are mapped to major lineages using a dataset-specific keyword map (Table S3), and concordance is assessed in a descendant-aware manner: a fine-grained label counts as concordant with a coarser reference label, and vice versa, when their lineage sets overlap [8].

Cytometry has no universally adopted cell-type nomenclature, so the same population may be named differently by different studies, tools and annotators [9]. Scoring an annotation against a reference therefore requires deciding when two differently worded labels refer to the same population. The keyword map reduces free-text labels to a small set of major lineages so that reference, model and expert annotations can be compared on equal terms. It is used only for scoring. Ontology-based tools address the separate problem of deriving standardised names for populations from their marker profiles [9, 3].

Because annotators and models may describe the same population in different words, this map is applied identically to reference labels, expert free-text annotations and model output. Without it, the comparison would measure vocabulary differences instead of cell identity. The maps were completed iteratively: any label appearing unmapped was resolved to its major lineage and the corresponding synonym was added. Every entry is an objectively correct lineage assignment and is auditable in Table S3, and unrecognised labels are scored as disagreements and not excluded. Across all model–dataset combinations with an unmapped fraction above 2%, every distinct unmapped label was a coarse term spanning several lineages, an explicit abstention, or a population the reference itself never gated; none was a specific single-lineage cell type that the map failed to cover. Unmapped fractions track model behaviour rather than map coverage: the highest fraction is observed for the frontier control Opus 5 (39.1% on Samusik2016), while Claude Sonnet has the highest unmapped fraction among the three strongest models on Parrish2023. Per model and dataset values are given in Table S4.

#### Unit of analysis

The LLM makes a single annotation decision for each metacluster, which is then inherited by all constituent clusters. Treating individual clusters as independent observations would therefore pseudo-replicate each decision *n*_cluster_*/n*_metacluster_ times and inflate apparent precision. The metacluster is therefore the primary unit of evaluation: its reference lineage set is the union of those of its constituent clusters, and concordance is the frequency-weighted proportion of metaclusters for which the LLM and reference lineage sets overlap. Unweighted concordance was also computed; it showed the same pattern and did not alter the model ranking.

#### Uncertainty

95% confidence intervals are reported for every metacluster-level concordance estimate, obtained by nonparametric bootstrap with 2,000 resamples of the metaclusters drawn with replacement and percentile-based intervals. For comparisons between two scoring methods on the same data (§3.4) a paired bootstrap is used: each resample is shared across both methods, allowing the difference Δ to be calculated within each resample. The 95% confidence interval for Δ is then defined by the 2.5th and 97.5th percentiles of its bootstrap distribution, and an interval excluding zero is interpreted as a statistically significant difference.

#### Chance correction

Raw concordance cannot distinguish a model that resolves fine-grained cell identity from one that hedges to a coarse, multi-lineage label that receives credit for every lineage it covers, nor can it be interpreted meaningfully without comparison to a trivial baseline. Three complementary measures are therefore reported for each model: (i) a majority-class baseline, obtained by always predicting the most frequent reference lineage (Cohen’s *κ* = 0 by construction); (ii) metacluster-level Cohen’s *κ*, interpreted using the conventional Landis–Koch scale [20]; and (iii) the coarse-label fraction, defined as the frequency-weighted proportion of cells assigned a label spanning more than one lineage. Cohen’s *κ* is computed over single categorical labels, using the frequency-weighted majority reference lineage and the model’s display lineage for each metacluster. Multi-lineage model labels are treated as distinct categorical outcomes and do not receive credit through set overlap. The descendant-aware matching used for concordance is therefore deliberately not applied to *κ*, making it the stricter metric.

#### Composition

Major-lineage proportions are compared via the total-variation distance (TVD) between reference and model proportion vectors, defined as half the sum of absolute differences reported in percentage points. The comparison is restricted to clusters for which both the reference and the model commit to a single lineage. Composition is a cell-level quantity, so unlike concordance and *κ*, it is computed per cluster, not per metacluster. TVD is a single distributional summary rather than a per-unit mean, so the pseudo-replication concern described above does not apply. Between 89% and 99% of clusters are retained across datasets. Pearson/Spearman correlations are not used because they can be dominated by the wide dynamic range of lineage proportions.

#### Method-matched inter-expert ceiling

For Capelle2023, two experts (H2, H3) independently annotated the same 40 metaclusters provided to the model, from the continuous heatmap alone. Each expert used their own terminology and received no population-definition sheet. Both were blinded to the CyCadas reference (H1), to each other’s annotations, and to all model outputs. Because H2 and H3 received the same information as the models, their agreement provides a method-matched ceiling. Their free-text labels were reconciled to lineages and fine attributes using the same mapping rules applied to the models. All 40 metaclusters could be mapped for both annotators without extending the mapping, using the same final map applied to the models. Both annotators are co-authors but contributed to neither CyCadas nor the LLM pipeline, making this ceiling independent of the method being evaluated. Four annotations are compared (H2, H3, the binary-gated H1, and each LLM) at the metacluster level using descendant-aware concordance, both for major lineages and for fine attributes (T coreceptor, maturation state, monocyte subset; §3.3).

#### Hand-curated marker-matrix comparator

For Capelle2023 only, a deterministic, LLM-free marker-matching rule is also reported, scored against a reference matrix built once, by hand, from the target-population vocabulary (§3.1). It serves two purposes: illustrating the limitations of raw concordance, which can reward hedging to coarse labels, and providing a meaningful baseline for comparison for the LLM-based approach. Robustness of the hand-curated marker definitions was assessed by single-marker perturbation: 214 edits, comprising sign reversals and marker removals, were applied individually, with *κ* recomputed after each edit.

#### Input-representation controls

Two controls test whether the results depend on how profiles are presented. In a serialisation ablation, the three-band encoding was compared with the raw scaled marker values and with a binary positive/negative representation, using Claude Sonnet on Capelle2023, Samusik2016 and Levine2015, with the same metacluster partition and annotation procedure across conditions. In a marker-name permutation control, marker labels were randomly permuted using a fixed random seed, so that each intensity vector was presented under the incorrect marker name. The model was then rerun and evaluated against the true reference. Because permutation destroys the biological meaning of the profiles, metacluster structure is no longer interpretable; both controls are therefore reported per-cluster (§3.6).

#### Intermediate-expression analysis

Because binary gating cannot represent intermediate expression levels, whereas the direct method can, disagreements between the two approaches are expected to concentrate on metaclusters containing more markers with intermediate expression. This was tested using the intermediate-band fraction, defined as the proportion of marker values falling within the symmetric band [0.35, 0.65] after scaling. This interval was chosen independently of the ≥ 0.6/*<* 0.35 bands used to serialise the prompt. Disagreeing and agreeing metaclusters were compared using a one-sided Mann–Whitney *U* test (alternative: disagreeing *>* agreeing). The comparison was made at the metacluster level, since per-cluster *p*-values are inflated by pseudo-replication, and used Claude’s disagreements, which are least likely to reflect model error. *p*-values are not corrected for multiple comparisons.

Concordance stratified by self-reported confidence and reproducibility across runs for all datasets are also reported.

Majority-vote consensus was computed at the metacluster level by taking the most frequent major-lineage assignment across the seven models, then scoring it against the reference using the same descendant-aware rule applied to individual models. Where two or more lineages received the same highest number of votes, the metacluster was scored as a disagreement, since a tied vote produces no consensus label. Breaking such ties in favour of one model would make the result depend on the order in which model outputs were read, and would allow the consensus label to inherit the prediction of the model it is being compared against. Ties occurred only on Samusik2016 (5 of 40 metaclusters, 4.3% of cells); Levine2015 and Parrish2023 had none.

### 2.7 Implementation

The annotation method itself is deliberately compact: a single routine scales the cluster medians, forms metaclusters, serialises each profile, issues the LLM call, and propagates the returned labels. The implementation uses standard scientific Python libraries and supports interchangeable cloud (Anthropic) and local (OpenAI-compatible/Ollama) LLM backends. The same engine is wrapped in a lightweight interactive application in which users upload a cluster table and obtain cell-type labels, representing the intended mode of routine use. The remainder of the codebase is devoted to evaluation, including dataset preparation, concordance, *κ*, and human-agreement-ceiling analyses, robustness and reproducibility assessments, and figure generation. All analyses are executable from the command line and are documented in the repository. Source code and a reading guide to the scripts are available at the PhenoScribe repository: https://git.lih.lu/bioinformatics-and-ai/phenoscribe (release v1.0.1, MIT licence), archived on Zenodo [21].

## 3 Results

### 3.1 LLM-based annotation reproduces expert annotation

The LLM-based method assigned a label to 100% of clusters. Fig. 2 illustrates the annotation of the Capelle2023 Parkinson’s disease whole-blood dataset, showing reference and model assignments alongside the corresponding marker profiles. Every cluster received a label, although some models return a coarse or “Undefined” label instead of a specific call (§3.4). Because the LLM makes one decision per metacluster, performance was evaluated at the metacluster level (Table 2, Fig. 3). A single well-chosen model reaches 85–98% metacluster concordance across all four datasets. This is practically relevant since a model must be selected before any reference annotation exists to check it against (cloud Claude Sonnet 85.2–98.2%; the best local model, qwen3.6, 83.3–96.0%). The trivial majority-class baselines (44.6–64.0%, *κ* = 0) are cleared by a wide margin on chance-corrected agreement (Table 3). Composition error, measured as the TVD between the lineage-proportion vectors of the best-performing model and the reference, is low on most datasets: 0.03 percentage points for Capelle2023, where the model’s lineage proportions almost exactly reproduce the reference, 3.5 for Samusik2016 and 7.4 for Levine2015. The largest difference is on Parrish2023 (8.5 points). TVD is reported for the highest-concordance model on each dataset: Claude Sonnet for Capelle2023, Samusik2016 and Parrish2023, and Llama 4 for Levine2015; on Parrish2023 the three leading models are separated by 0.2 percentage points, so that designation is nominal. TVD can exceed the nominal 100 − concordance bound because descendant-aware scoring considers a metacluster concordant when the model lineage is contained within the reference lineage set, including multi-lineage reference labels. A metacluster can therefore be scored concordant while still shifting mass between lineages in the proportion vector, making TVD the stricter composition check. Because raw concordance can reward hedging to coarse labels, it is interpreted together with Cohen’s *κ* and the coarse-label fraction.

**Figure 2:**
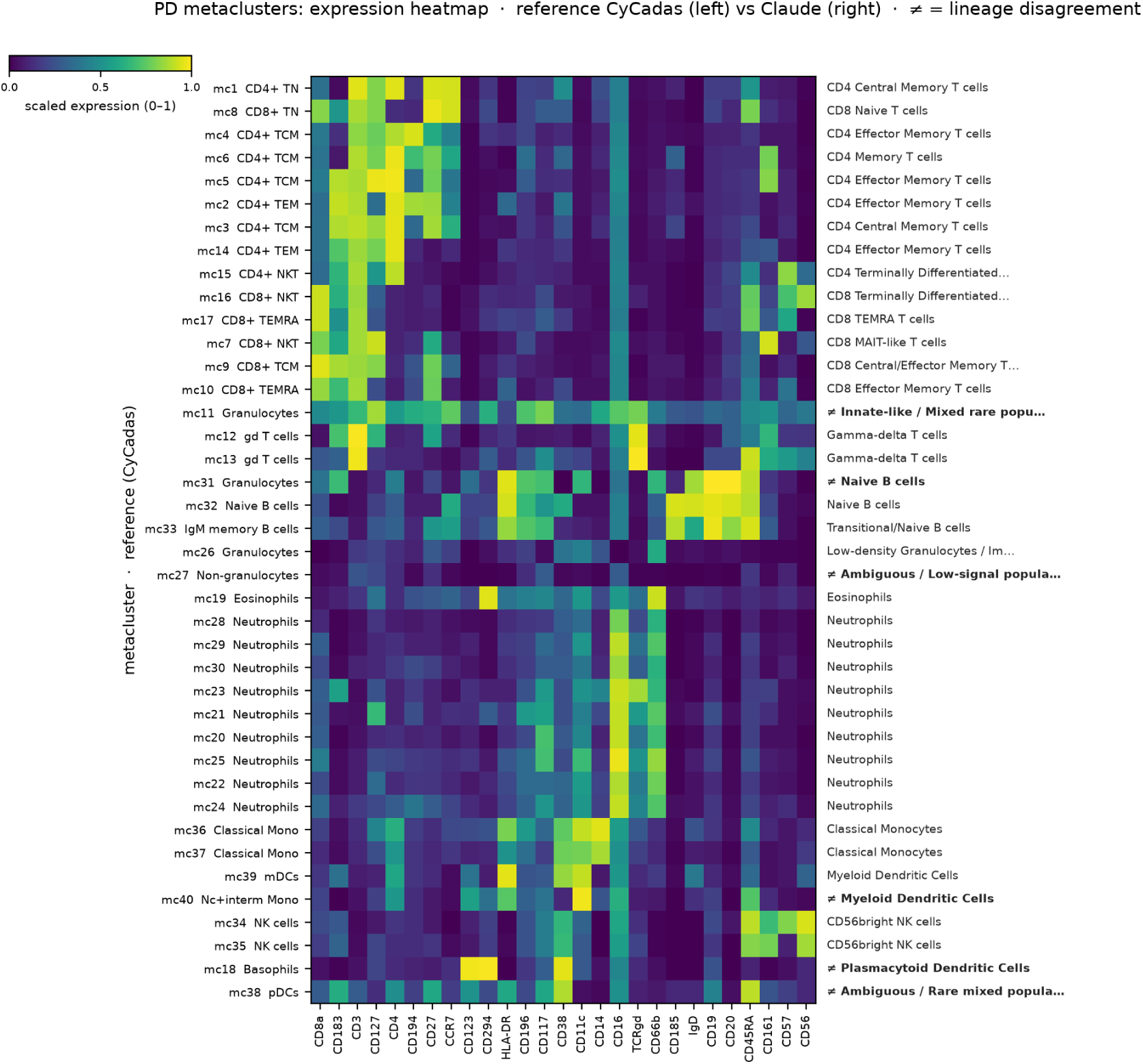
Capelle2023 metacluster annotation, reference vs. model. The 40 Capelle2023 metaclusters (rows) × 27 markers (columns), scaled 0–1 expression (viridis, sequential), Ward-clustered on both axes. Each row’s reference cell type (H1) is the left-hand label and the Claude Sonnet label is on the right; lineage-level disagreements are marked ̸=. Only 6/40 metaclusters disagree, and they fall on established boundaries (basophil vs. pDC, monocyte vs. DC, and coarse “Granulocyte”/“Non-granulocyte” reference buckets), not gross errors.

**Figure 3:**
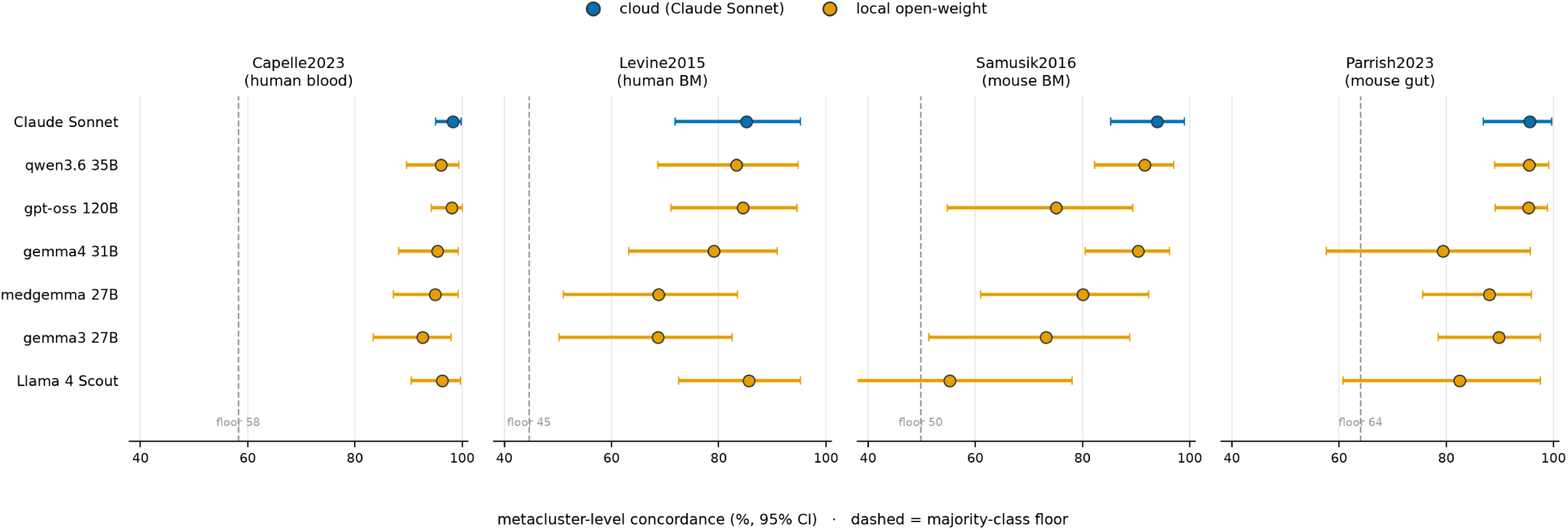
Metacluster-level concordance across seven models and four datasets, with 95% bootstrap CIs. Each panel is one dataset; models are ordered by mean concordance (identity by row position). Points are frequency-weighted metacluster concordance, whiskers the 95% bootstrap CI, and the dashed line the majority-class floor. Colour distinguishes the cloud model (Claude Sonnet) from the six local open-weight models (Okabe–Ito, CVD-safe). Every model outperforms its corresponding baseline, but the ranking reshuffles across domains, with no universal best *local* model (Llama 4 is comparable to the top models on Levine2015 yet collapses on Samusik2016; gpt-oss is top-tier on Capelle2023/Parrish2023 yet drops on Samusik2016), while the single cloud model stays near the top throughout.

**Table 2:**
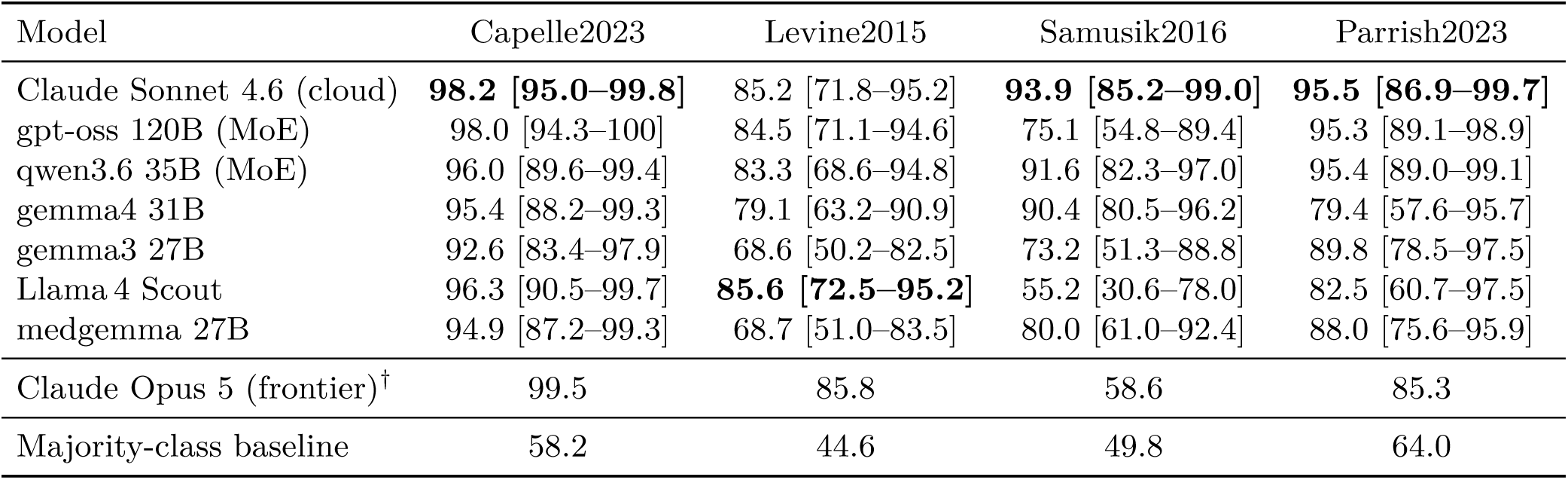
Metacluster-level concordance (%, frequency-weighted, 95% bootstrap CI) across all four datasets. This is the correct unit, since the LLM makes one decision per metacluster. Metaclusters: 40 per dataset (Samusik2016 and Parrish2023 are mouse). Bold indicates the best result for each dataset. The majority-class baseline represents the performance floor obtained by always predicting the most frequent lineage. No model consistently performs best across datasets and the ranking reshuffles by domain (Llama 4 comparable on Levine2015, worst on Samusik2016). Models as of July 2026. **^ψ^**Claude Opus 5 is included as a frontier control, not among the seven evaluated models. Because the API does not accept a temperature parameter for this model, its outputs are non-deterministic; reported values are therefore the mean of three runs, with the range given in §3.5. Its low Samusik2016 score reflects a reproducible lineage error (§3.4).

**Table 3:**
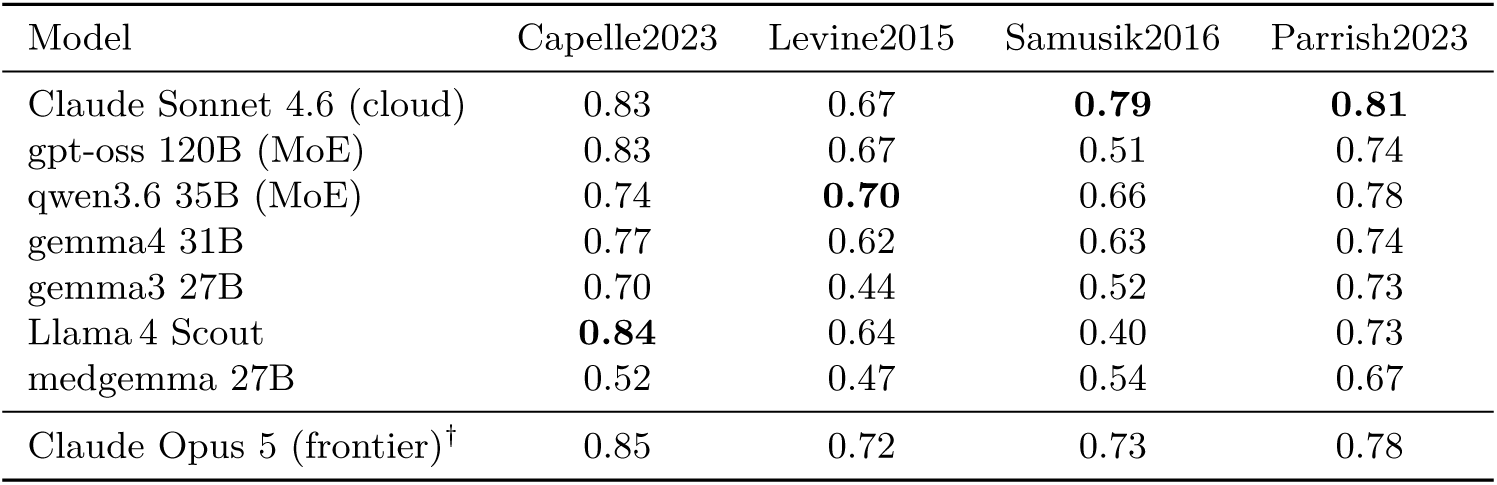
Chance-corrected agreement (metacluster-level Cohen’s *κ*_mc_) across all four datasets; the majority-class baseline is *κ* = 0 by construction. Every model performs above chance on every dataset, but performance varies substantially: weaker models, such as Llama 4 on Samusik2016 and gemma3/medgemma on the bone-marrow benchmarks, show marked drops, and no model consistently performs best across all datasets. Hedging is rare, except medgemma on Capelle2023, where 60.7% of cells receive a coarse *>*1-lineage label (§3.4). Bold indicates the best result for each dataset. **^ψ^**Frontier control (Opus 5), reported as the mean over three runs. Its Samusik2016 *κ* remains substantial despite a marked drop in concordance (Table 2) because *κ* is computed with equal weight across metaclusters whereas concordance is cell-weighted, and Opus 5’s errors are concentrated in a small number of large metaclusters.

| Model | Capelle2023 | Levine2015 | Samusik2016 | Parrish2023 |
| --- | --- | --- | --- | --- |
| Claude Sonnet 4.6 (cloud) | 0.83 | 0.67 | **0.79** | **0.81** |
| gpt-oss 120B (MoE) | 0.83 | 0.67 | 0.51 | 0.74 |
| qwen3.6 35B (MoE) | 0.74 | **0.70** | 0.66 | 0.78 |
| gemma4 31B | 0.77 | 0.62 | 0.63 | 0.74 |
| gemma3 27B | 0.70 | 0.44 | 0.52 | 0.73 |
| Llama 4 Scout | **0.84** | 0.64 | 0.40 | 0.73 |
| medgemma 27B | 0.52 | 0.47 | 0.54 | 0.67 |
| Claude Opus 5 (frontier)<sup>†</sup> | 0.85 | 0.72 | 0.73 | 0.78 |

A deterministic, LLM-free marker-matching rule based on a hand-built Capelle2023 reference matrix shows the effect of coarse-label fallback directly. When fallback is allowed, the rule uses it for 35.3% of cells and achieves a chance-corrected agreement of *κ* = 0.40; when constrained to make a specific cell-type assignment, *κ* increases to 0.879. Raw concordance is therefore strongly affected by whether coarse-label fallback is allowed, as also observed for MedGemma (§3.4). On the panel for which it was developed, the matrix matches or exceeds the performance of every LLM tested. The rule is also robust to perturbation: among the 214 single-marker edits, only modifications to CD56 in the NK-cell definition substantially reduced *κ* (from 0.88 to 0.19–0.24); the remaining 212 edits had little effect.

Model-reported confidence was associated with annotation concordance, although formal calibration was not assessed (no reliability diagram was constructed). On Capelle2023, high-confidence predictions showed 94–98% concordance across models (Claude 96.4%, gpt-oss 98.3%, qwen3.6 94.1%), whereas concordance in the low-confidence category was substantially lower (Claude 20%, qwen3.6 0%; approximately 20 metaclusters per model). The relationship was not strictly monotonic: for several models, medium- and high-confidence predictions had similar concordance, while one model achieved 100% concordance in its low-confidence category based on only eight metaclusters. Reported confidence therefore provides a useful discriminative signal for flagging annotations that warrant human review, but should not be interpreted as a calibrated probability of correctness. Most disagreements occurred between biologically related populations, including basophils versus plasmacytoid dendritic cells (pDC), both of which express CD123, and monocytes versus dendritic cells (DC).

### 3.2 Concordance generalises across annotation paradigms and species

Capelle2023 was annotated using CyCadas, whereas the remaining three datasets represent different annotation paradigms. Levine2015 and Samusik2016 are public benchmarks with manual-gating references from independent studies, whereas Samusik2016 additionally provides a cross-species evaluation in mouse. **Parrish2023** was annotated by direct expert inspection of the marker heatmap without a gating tool.

High concordance was observed across all three datasets (Table 2). On Levine2015, the best models reached approximately 85% concordance, compared with a majority-class baseline of 44.6%. On Samusik2016, concordance reached 93.9% for Claude Sonnet, 91.6% for qwen3.6 and 90.4% for gemma4, against a baseline of 49.8%. On Parrish2023, whose reference is the paradigm closest to the LLM’s own task, the best models achieved approximately 95% (Claude Sonnet 95.5%, qwen3.6 95.4%, gpt-oss 95.3%), against a baseline of 64.0%. Concordance with a direct-heatmap human reference is therefore comparable to that with the threshold-based CyCadas reference, across both human and mouse datasets.

Metacluster purity remained high across all three datasets (99.6–100%), indicating little major-lineage mixing within metaclusters. The concordance therefore reflects genuine labelling agreement and is not an artefact of the clustering.

Two further observations are noteworthy. Every model, including the cloud model, assigned a block of reference- “Monocyte” clusters in Levine2015 to DC, progenitor or megakaryocyte populations. These clusters are CD41^+^ CD61^+^CD133^+^, consistent with platelet and progenitor signatures. This accounts for most of the gap between bone marrow (∼85%) and blood (∼96%) concordance. Part of that gap is structural: the Levine2015 reference contains no conventional dendritic cell, neutrophil or megakaryocyte population, while Samusik2016 contains no neutrophil or ILC population. Consequently, models that correctly identify these populations cannot be mapped to a reference lineage and are scored as disagreements.

### 3.3 Models fall within the method-matched inter-expert ceiling

Using the method-matched inter-expert comparison described in §2.6, H2 and H3 agreed on 99.3% of metaclusters at the major-lineage level (frequency-weighted). Claude Sonnet and gpt-oss showed similarly high concordance, both at approximately 99%. Because agreement at the lineage level was nearly complete for both humans and models, this level provided limited discrimination.

Differences became apparent at finer annotation levels (Table 4). CD4/CD8 coreceptor assignment showed complete expert agreement (100%), with the best-performing models also approaching 100%. For T-cell maturation state (TN, TCM, TEM and TEMRA), however, H2 and H3 agreed on 73.3% of metaclusters (*n* = 15; bootstrap 95% CI 47–93%). Model concordance with the two experts ranged from 64% to 88%: Claude Sonnet agreed with H2 and H3 on 77% and 69% of metaclusters respectively, gpt-oss on 88% and 75%, and qwen3.6 on 71% and 64%. This range brackets the 73.3% at which the two experts agree. The binary-gated reference (H1) showed similar concordance with the two continuous-profile experts, agreeing with H2 on 67% and with H3 on 75% of maturation-state assignments, so the lower agreement for maturation state was also present when comparing a binary-gated reference with independent continuous-profile annotations.

**Table 4:**
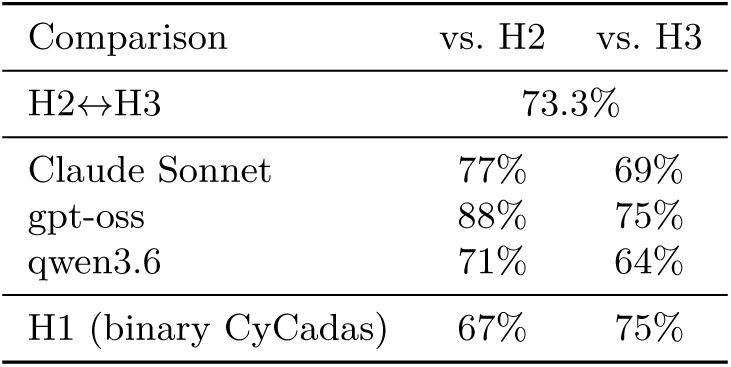
Agreement on T-cell maturation state (TN/TCM/TEM/TEMRA) for Capelle2023, at the metacluster level and descendant-aware. *n*=15 metaclusters where both annotators commit a value; bootstrap 95% CI for the expert pair 47–93%. H2 and H3 each annotated from the continuous heatmap alone; H1 is the binary-gated CyCadas reference. Agreement at the major-lineage and coreceptor levels is near-complete for all annotators and models and is given in the text.

| Comparison | vs. H2 | vs. H3 |
| --- | --- | --- |
| H2↔H3 | 73.3% |  |
| Claude Sonnet | 77% | 69% |
| gpt-oss | 88% | 75% |
| qwen3.6 | 71% | 64% |
| H1 (binary CyCadas) | 67% | 75% |

This estimate is based on a single pair of annotators and 15 maturation-state metaclusters, giving a wide confidence interval.

### 3.4 Model reliability is dataset-dependent

Model performance varied substantially across datasets (Table 2, Fig. 3). Capelle2023 showed little separation between models, with overlapping confidence intervals and concordance approaching the ∼99% lineage-purity limit. In contrast, the two bone-marrow benchmarks and the Parrish2023 gut panel showed clear differences between models.

On Samusik2016, Claude Sonnet achieved the highest concordance (93.9%), followed by qwen3.6 (91.6%) and gemma4 (90.4%), whereas Llama 4 showed substantially lower concordance (55.2%). On Parrish2023, the three leading models were separated by 0.2 percentage points (Claude Sonnet 95.5%, qwen3.6 95.4%, gpt-oss 95.3%), far below the width of their confidence intervals, so no ordering among them is claimed; gemma4 showed lower concordance (79.4%). Complete results for all models and datasets are given in Table 2.

Model rankings were not consistent across datasets. Llama 4 performed among the stronger models on Levine2015 but was the lowest-performing model on Samusik2016, while gpt-oss performed among the strongest on Capelle2023 and Parrish2023 but showed lower concordance on Samusik2016 (75.1%). No local model ranked first across all four datasets. Claude Sonnet achieved at least 85% concordance on every dataset and was best or second-best on all four, ranking first on three, although on Parrish2023 its lead over qwen3.6 and gpt-oss falls within the corresponding confidence intervals.

The models also differed in their label-assignment behaviour. On Samusik2016, 22.5% of Llama 4’s labels were unmapped to the reference vocabulary (“Myeloid Cells”, “Dendritic Cells”, and non-reference Erythroblasts and Megakaryocytes). On Parrish2023, gemma4 assigned “Undefined” to 14% of cells while retaining a *κ* of 0.74.

#### Frontier LLM is not a shortcut to reliability

Claude Opus 5 showed dataset-dependent performance similar to the other models (Table 2). On Capelle2023 it achieved 99.5% concordance, and on Levine2015 85.8%. On Samusik2016, however, Opus 5 achieved only 58.6%, substantially below Claude Sonnet and the strongest local models. On Parrish2023 it reached 85.3% (mean of three runs, range 81.4–92.0), also below the strongest models, and produced a higher proportion of “undefined / lineage-negative” assignments.

The Samusik2016 result was consistent across all three runs. In each, Opus 5 assigned five large Ly6C^hi^CD11b^+^ metaclusters to neutrophils or granulocyte precursors, whereas the manual-gating reference, Claude Sonnet and three local models assigned them to monocytes. The neutrophil marker Ly6G is low in exactly these metaclusters (≤0.26 on the 0–1 scale). These five metaclusters represent 38% of cells, producing a substantial impact on cell-weighted concordance, while the unweighted metacluster-level *κ* remains substantial (0.73). Much of this reflects off-vocabulary output, not misassignment within the reference vocabulary: 39.1% of Opus 5’s cells on Samusik2016 were unmapped (Table S4), and that reference contains no neutrophil or granulocyte-precursor population, so these labels cannot map to a reference lineage. The concordance figure is unaffected either way, since the reference assigns these metaclusters to monocytes.

Majority-vote consensus was evaluated at the metacluster level on Levine2015, Samusik2016 and Parrish2023. On Levine2015, consensus achieved 89.3% concordance, exceeding the best individual model (Llama 4, 85.6%), though the difference was not significant (Δ = +3.7, 95% CI [−9.6, +17.3]). On Samusik2016, consensus achieved 84.6% concordance, significantly below Claude Sonnet (93.9%; Δ = −9.3, CI [−19.6, −3.1]). This was also the only dataset on which the models produced tied consensus labels (5 of 40 metaclusters, 4.3% of cells), and those metaclusters are scored as non-concordant. On Parrish2023 consensus (93.4%) was also lower than the best individual model, though not significantly (Δ = −2.1, CI [−12.5, +7.7]).

Model agreement was associated with reference concordance across all four datasets. Metaclusters for which all models produced the same lineage assignment were 99.3–100% concordant with the reference and contained 35–70% of cells, depending on the dataset. On the remaining split metaclusters, majority-vote concordance was 66–94% (both frequency-weighted, as in Table 2). The gap is smallest on Capelle2023 (99.3% versus 93.6%), where all models perform close to the purity limit.

Medical fine-tuning did not improve performance over the corresponding base model. On Capelle2023, medgemma produced coarse labels for 60.7% of cells. On Levine2015 its concordance was 68.7%, essentially identical to gemma3 (68.6%). Full comparisons are given in Table 2.

### 3.5 Reproducibility

Run-to-run reproducibility was high for the stronger models at the major-lineage level. Lineage reproducibility is the share of cells whose metacluster received the same major lineage in all three runs; a label that maps to no reference lineage is treated as its own category. qwen3.6 produced identical lineage assignments for at least 96.8% of cells on every dataset, while Claude Sonnet showed 82–99.8% lineage agreement across runs. Exact fine-label agreement was lower, reflecting variation in label wording despite greater stability at the lineage level. gemma3 showed greater dataset-dependent variation. On Capelle2023, only 40% of cells received the same lineage assignment across runs, driven primarily by changes in the assignment of the large granulocyte-associated metaclusters between “Neutrophil” and “Monocyte”. On Samusik2016, lineage agreement reached 85%, whereas it was 100% on the other two datasets.

Claude Opus 5 showed high lineage reproducibility on Capelle2023 (100%), Samusik2016 (96.5%) and Levine2015 (94.8%), and lower agreement on Parrish2023 (84.0%). On Samusik2016, the five metaclusters mislabelled as granulocytic (§3.4) received the same incorrect lineage in all three runs, with only the terminology varying between terms such as “mature neutrophils”, “myelocytes” and “granulocytic precursors”. Thus, the underlying error was reproducible even though the label string was not.

### 3.6 Labels depend on marker identity, not on the serialisation

Permuting marker names substantially reduced annotation concordance on Capelle2023. With the original marker identities, per-cluster concordance was 95.8% for gpt-oss and 95.5% for Claude Sonnet. After randomly permuting the marker names while preserving the underlying intensity values, concordance fell to 15.0% and 24.4%, respectively. Both values are well below the per-cluster majority-class baseline of 53.2%, so the labels depend on which marker each value belongs to, not on the pattern of values alone.

The serialisation itself had little effect. Replacing the three-band encoding with the raw scaled marker values changed per-cluster concordance by +0.0, +1.0 and −1.0 percentage points on Capelle2023, Samusik2016 and Levine2015 respectively, whereas reducing the profiles to binary positive/negative calls changed it by +0.1, −9.0 and +8.0 points. Graded representations, three-band or raw, therefore gave similar results, whereas binary encoding produced substantially larger changes on two of the three datasets.

### 3.7 Disagreements track intermediate expression

Disagreeing metaclusters contained a higher fraction of intermediate-range marker expression than agreeing meta-clusters on both manual-gating benchmarks. On Samusik2016 the intermediate-expression fraction was 1.67-fold higher among disagreeing metaclusters (*p* = 0.002), and on Levine2015 1.47-fold higher (*p* = 0.028). The same pattern was observed on Samusik2016 with qwen3.6 (*p* = 0.016). The enrichment was weak or absent on the saturated Capelle2023 blood dataset, and could not be reliably assessed on Parrish2023 because too few metaclusters were discordant.

## 4 Discussion

Our central methodological finding is that an LLM can annotate cell populations directly from continuous cluster-level expression profiles, using the same type of information available to an expert reading a heatmap. The approach requires no gating tree, hand-authored per-marker thresholds, training data, or panel-specific classifier, and produces annotations in a single LLM call. Because the model is given only a serialised list of scaled marker values, it is reasonable to ask whether the resulting annotations depend on the biological identity of the markers or merely on the numerical pattern of their expression values. The marker-name permutation experiment indicates that marker identity does matter: when marker names were randomly permuted while preserving the underlying expression values, concordance dropped sharply for both gpt-oss and Claude Sonnet, from approximately 96% to 15–24% (§3.6). The model therefore does not assign labels from the numerical pattern of expression values independently of marker identity.

A key challenge in evaluating cell-type annotation is the absence of a definitive ground truth. Manual gating is itself a human interpretation and is not perfectly reproducible between experts. We therefore do not interpret concordance with any single reference as absolute accuracy. Instead, we estimated a method-matched human ceiling using two independent expert annotations of the same continuous profiles available to the LLM (§3.3). At the major-lineage level the two experts showed near-complete agreement, leaving little room to distinguish methods. On T-cell maturation state they agreed less strongly, at 73.3%. The strongest LLMs agreed with the two experts across a band that brackets this value, so disagreement between the LLM and an individual expert does not necessarily indicate an incorrect annotation, given the level of disagreement between independent human readers.

This is particularly relevant for intermediate expression. Disagreements with the binary-gated references were enriched for intermediate-range marker expression on both manual-gating benchmarks, with the same pattern reproduced across models on Samusik2016 (§3.7). Some LLM–reference differences may therefore arise because the LLM uses graded expression information that is not represented in binary gating, which would be most relevant for populations with overlapping marker profiles or gradual biological transitions. Consistent with this, collapsing the input itself to binary calls destabilises the annotation (§3.6). However, the enrichment alone does not establish that the LLM’s alternative annotation is more accurate; it indicates only that intermediate expression is associated with cases in which the LLM and the binary reference disagree. The same principle is illustrated by the Levine2015 monocyte-to-progenitor disagreement: every model tested assigned reference-“Monocyte” clusters to DC, progenitor or megakaryocyte populations, and these clusters showed CD41, CD61 and CD133 expression consistent with platelet- and progenitor-associated profiles. Such cases highlight the difficulty of treating a single categorical reference as an error-free ground truth when biologically heterogeneous or transitional populations are involved.

The ceiling also indicates how concordance should be read. Judged against perfect agreement, a value of 85% appears weak; judged against what two experts achieve on the same task, it is close to the attainable limit at the major-lineage level and above it at finer granularity. Concordance should therefore be interpreted relative to the agreement achievable between human annotators.

Generalisation across annotation tools, paradigms and species is a further finding. The method generalised beyond the CyCadas dataset to three independent datasets: Levine2015 and Samusik2016 use manual-gating references from independent studies, with Samusik2016 additionally representing a different species, and Parrish2023 was annotated by direct expert inspection of a marker heatmap rather than by a gating tool. Concordance remained high across all three datasets, ranging from 85% on Levine2015 to 95% on Parrish2023. These results reduce the possibility that the observed concordance is driven primarily by reproducing the conventions of a single annotation tool. The Parrish2023 result is particularly informative because its reference was generated by reading marker patterns directly from a heatmap, closely matching the information available to the LLM. High concordance across these distinct reference types therefore supports the generalisability of the approach, while not implying that any single annotation paradigm provides a definitive ground truth.

LLM-based annotation is nonetheless model- and domain-dependent, with no universally best local model across panels and species. Llama 4 performed competitively on human bone marrow but showed substantially lower concordance on mouse bone marrow, while Claude Sonnet was the most consistent model in concordance across datasets. Greater model capability did not necessarily translate into greater reliability: Claude Opus 5 repeatedly assigned several large Ly6C^hi^CD11b^+^ mouse bone-marrow metaclusters to granulocytic populations despite low Ly6G expression, and the same assignment occurred across all three runs, indicating a reproducible rather than stochastic error (§3.4). Model scale alone therefore cannot substitute for validation on the target panel and species. Models also exhibited distinct failure modes. Llama 4 tended to produce coarse or off-vocabulary labels on Samusik2016, whereas gemma4 more often abstained through “Undefined” assignments on Parrish2023. These behaviours have different implications for quality control: abstention provides an explicit review signal, whereas an inappropriate but specific label may be harder to detect. Reproducibility likewise differed between models and datasets, with strong models showing stable lineage assignments even when fine-label wording varied, while weaker models showed substantial run-to-run lineage variation on difficult datasets. Model consensus did not consistently improve performance: majority voting increased concordance on Levine2015, though not significantly, but significantly reduced it on Samusik2016, where several models shared a similar failure mode. Consensus is therefore beneficial when model errors differ but can reinforce them when multiple models make the same incorrect assignment. In practice the expert reviews the returned labels instead of producing them. Because no model ranked first everywhere and rankings shifted across datasets, a model that performs well on one panel should not be assumed to transfer to another. Three signals in the output indicate where review effort should go. First, abstentions and low reported confidence flag uncertain calls. Second, the returned lineage proportions can be compared against expectation for the panel, which catches composition-level errors that per-cluster agreement can miss. Third, metaclusters with a high fraction of intermediate-range marker expression are where disagreement concentrates (§3.7).

Local open-weight models were competitive with Claude Sonnet on the independent benchmarks, demonstrating that the method can be deployed without a cloud service. This may be important for clinical or patient-derived data subject to restrictions on external processing. Although the method sends only aggregate per-marker expression profiles rather than single-cell or participant-level data to the LLM, these data may still be subject to institutional data-governance requirements. Local deployment can therefore provide an additional safeguard by keeping the data within the institution rather than transmitting them to an external service.

The closest prior work applies LLMs to single-cell RNA-seq cluster annotation from marker-gene lists [7, 8]. The present results extend that setting to continuous protein intensities, where the input is a graded profile rather than a discrete gene list and the reference is itself a human reading of the same profile. Relative to deep-learning annotators [4, 5, 6], the approach needs no training data, labelled examples or explicit panel encoding, and returns human-readable hierarchical labels. The hand-curated marker matrix matches or exceeds the LLMs on the panel it was built for, and is robust to all but one marker definition. Its cost is therefore curation rather than accuracy or fragility: the matrix must be built by hand for each panel and tissue, and applying it to the bone-marrow (32/39 markers) or mouse-gut panels would require rewriting it from scratch. The LLM removes that burden, which is why the matrix serves as a metric-design illustration on its home panel rather than a method the LLM must beat elsewhere.

### Limitations

First, one of the four references, Capelle2023, was generated using CyCadas, a tool one author co-developed, so this comparison may partly reflect a shared annotation framework; cross-lab generalisation therefore rests primarily on Levine2015 and Samusik2016, while Parrish2023 provides a cross-paradigm test based on direct heatmap inspection. The method-matched expert comparison adds an independent benchmark, but only for Capelle2023. Second, fine clustering of the bone-marrow datasets used *k*-means rather than a self-organising-map method; cluster purity was high (96–100%) and the base clustering already varies across the four datasets, which we read as robustness rather than a confound, but the two methods were not compared directly on the same dataset. Third, the inter-expert ceiling rests on two annotators and one dataset, with few metaclusters for some fine-grained attributes, and is indicative rather than definitive. Fourth, the marker-name permutation control was run on two models and on Capelle2023 only, the hand-curated marker-matrix comparator was evaluated only on the panel for which it was built, and metacluster granularity was not systematically varied. Fifth, the local models were not served at matched numerical precision: qwen3.6 ran at bf16 while the remaining five used default distribution tags quantised to approximately 4 bits per parameter, so the local rankings may partly reflect precision rather than model capability. Sixth, all results are specific to the model versions evaluated (July 2026), and newer versions may perform differently.

Finally, concordance depends on the lineage map. The map is part of the evaluation, not of the method: it exists so that labels produced by different studies, tools and annotators can be scored against one another, and within a single study the returned labels can be inspected directly without it. Its influence is measured rather than assumed. Unmapped fractions are reported per model and dataset (Table S4), and re-scoring under a map restricted to reference-grounded keywords reduces concordance substantially (Parrish2023 from 95.5% to 71.9%, with unmapped fractions rising to 27.6–76.4%; Table S5), because such a map counts correct standard nomenclature as a disagreement whenever it differs from the reference’s shorthand. Those values are a lower bound from a deliberately unfavourable scorer, not a corrected estimate. The qualitative conclusions hold under both maps, but per-dataset orderings are coverage-sensitive: under the stricter map gpt-oss overtakes qwen3.6 on Parrish2023, so part of the ranking reflects vocabulary alignment with the reference and not cell-type identification alone. Labels naming two lineages outright, such as “RORgt+ ILCs/T cells”, are resolved by rule precedence, which costs gemma3 3.3 percentage points on one Parrish2023 metacluster whose reference itself splits across three lineages. Most importantly, concordance with an existing annotation does not establish correctness.

Broader transferability remains to be established across additional panels, tissues, species, annotators, clustering methods and annotation paradigms. What the present results show is agreement with several established references, and with independent human readers of the same continuous profiles, within the disagreement envelope those readers themselves exhibit.

## 5 Conclusion

LLM-based annotation of continuous cytometry profiles can achieve high concordance with existing expert annotations, well above chance-corrected baselines, without a predefined gating tree, hand-authored per-marker thresholds, or task-specific training data. The approach generalised across four datasets spanning different annotation tools, paradigms and species, and can be implemented with local open-weight models where data-protection requirements restrict external processing. Comparison with independent experts interpreting the same continuous marker-expression profiles showed that LLM–expert agreement can fall within the range of agreement observed between the experts themselves, particularly for fine-grained attributes where independent interpretations vary. These findings support LLM-based annotation as a flexible alternative to rule-based annotation, provided that model output is checked against the intended panel and species and that disagreement, confidence and lineage-proportion checks are used as review signals.

## Availability

Source code and scripts to reproduce the analyses: https://git.lih.lu/bioinformatics-and-ai/phenoscribe (release v1.0.1, MIT licence), archived on Zenodo [21]. Data availability: Levine2015 and Samusik2016 are available through the **HDCytoData** Bioconductor package [15], originally deposited in FlowRepository under FR-FCM-ZZPH. The Parrish2023 SPF subset is deposited on Zenodo under CC BY 4.0 [19], and the cluster medians and expert annotation table used here are distributed with that study’s analysis pipeline at https://github.com/DII-LIH-Luxembourg/amuc-allergy (v1.0). For Capelle2023, the FCS data are from [14] and the cluster annotation table is distributed as demonstration data with CyCadas [13]. The two expert annotation sheets (H2, H3) are included in the code repository.

## Acknowledgements

We thank the authors and depositors of the public datasets analysed here: Levine et al. [16] and Samusik et al. [17] (obtained via HDCytoData [15]), and Parrish et al. [18] (amuc SPF colonic data), for making their data available for reuse. We thank Sang-Yoon Kim for hosting the internal Ollama service used for the local-model experiments and for answering technical questions about it.

This research was funded in part by the Luxembourg Research Fund (FNR), grant reference “PRIDE21/ 16749720/NEXTIMMUNE2”. For the purpose of open access, the author has applied a Creative Commons Attribution 4.0 International (CC BY 4.0) license to any Author Accepted Manuscript version arising from this submission.

## Author contributions

O.H.: conceptualisation; methodology; software; formal analysis; investigation; data curation; visualisation; writing original draft; project administration. E.T.G.: investigation; resources; validation; data curation; writing, review and editing. T.-M.B.: investigation; validation; writing, review and editing. A.K.: writing, review and editing. V.D.: writing, review and editing. M.S.D.: investigation; resources; funding acquisition; writing, review and editing. P.V.N.: supervision; writing, review and editing.

## Ethics

No new human or animal data were collected for this study. All datasets analysed here were generated in previously published studies [14, 16, 17, 18], and the ethical approvals and consent arrangements for those studies are described in the corresponding publications. Levine2015 and Samusik2016 were obtained in de-identified form via HDCytoData [15].

## Use of generative AI

In addition to the models evaluated in this study, Claude (Anthropic) was used to assist with drafting and editing the manuscript text and with developing and running the analysis code. The authors reviewed and revised the AI-assisted text, tested and validated the code, and take full responsibility for the manuscript and the reported results.

