## Supplementary Tables S1-S5 for "Language models annotate mass cytometry clusters within the range where human experts disagree"

Table S3: Keyword maps used to reduce free-text labels to major lineages. Matching is case-insensitive on the label string; the first matching pattern determines the assigned lineage. Coarse groups (marked *) are matched before the lineage rules and map to a set of lineages (shown in parentheses); they are used only for the CyCadas dataset. Maps are given per dataset because each reference resolves to a different set of major lineages; the label-to-lineage relationships themselves are not panel-specific. Most patterns are matched as substrings, so "granulocyt" captures both "granulocyte" and "granulocytic"; patterns marked $^w$ are matched at word boundaries instead, without which "tem" would fire inside "Stem" and reassign haematopoietic stem cell labels to T. Patterns ending in a digit carry a negative lookahead for further digits, so "th1" does not match "th17". Rows are listed in the order the code tests them, and patterns within a row are tested left to right. "apc" maps to Myeloid because an unqualified antigen-presenting-cell label implies a non-B population, B cells being labelled as B cells. Fine attributes are applied only within the T-cell subset of the Capelle2023 inter-expert ceiling analysis and are not used for major-lineage scoring.

| Major lineage | Keyword patterns |
|---|---|
| **Capelle2023** (human whole blood) | |
| Non-granulocyte* | non-granulocyte, non granulocyte, nongranulocyte *(to T, B, NK, Monocyte, pDC, mDC)* |
| Granulocyte* | granulocyte *(to Neutrophil, Eosinophil, Basophil)* |
| DC* | dcs, dendritic cell, dendritic cells *(to pDC, mDC)* |
| Neutrophil | neutrophil |
| Eosinophil | eosinophil |
| Basophil | basophil |
| pDC | plasmacytoid, pdc |
| mDC | myeloid dendritic, conventional dendritic, mdc, cdc |
| B | b cell, b-cell, plasmablast, plasma cell, cd20 |
| Monocyte | monocyte, mono |
| T | nkt, mait, cd4, cd8, cd3, tcr, gamma, $\gamma\delta$, gd t, th1, th2, th17, treg, regulatory t, t cell, t-cell, temra, naive t, tcm$^w$, tem$^w$, trm$^w$, tfh$^w$, teff$^w$, th9$^w$, th22$^w$ |
| NK | nk, cd56, natural killer |
| **Levine2015** (human bone marrow) | |
| pDC | plasmacytoid, pdc |
| Basophil | basophil |
| NK | natural killer, nk, cd56 |
| HSPC | hspc, hsc, progenitor, stem cell, stem, cd34, cmp, gmp, mep, mpp, hematopoietic |
| Monocyte | monocyte, mono, cd14 |
| B | b cell, b-cell, b_cell, plasma cell, plasmablast, plasma, pre-b, pro-b, pre_b, pro_b, cd19, cd20 |
| T | t cell, t-cell, t_cell, thymocyte, tcr, cd4, cd8, cd3, naive t, tcm$^w$, tem$^w$, trm$^w$, tfh$^w$, teff$^w$, th9$^w$, th22$^w$ |
| **Samusik2016** (mouse bone marrow) | |
| pDC | plasmacytoid, pdc |
| mDC | myeloid dendritic, conventional dendritic, mdc, cdc |
| Basophil | basophil |

*continued on next page*

*Table S3 (continued)*

| Major lineage | Keyword patterns |
|---|---|
| Eosinophil | eosinophil |
| Mast | mast |
| Monocyte | monocyte, mono, macrophage, f4/80, f480 |
| T | cd4, cd8, cd3, tcrb, tcrgd, tcr, gd t, $\gamma\delta$, gamma delta, gamma-delta, nkt, mait, t cell, t-cell, thymocyte, treg, foxp3, t helper, helper t, th1, th2, th17, cytotoxic t, naive t, tcm$^w$, tem$^w$, trm$^w$, tfh$^w$, teff$^w$, th9$^w$, th22$^w$ |
| NK | natural killer, nk, nkp46 |
| HSPC | hspc$^w$, hsc$^w$, mpp$^w$, clp$^w$, cmp$^w$, gmp$^w$, mep$^w$, progenitor, stem cell, stem, multipotent, cd34, sca1, c-kit, ckit, hematopoietic, haematopoietic, lsk$^w$, myeloblast |
| B | b cell, b-cell, b_cell, b cells, b-cells, pro-b, pre-b, pro b, pre b, plasma, plasmablast, igd, igm, cd19, cd20, b220 |
| **Parrish2023 / amuc** (mouse colonic lamina propria) | |
| Mast | mast |
| Granulocyte | neutrophil, granulocyt, eosinophil, basophil |
| ILC | ilc, lti, innate lymphoid |
| T | th0, th1, th2, th17, treg, foxp3, cd8, cd4, dn t, dp t, t cell, t-cell, t lympho, cd3, tcr, nkt, gamma delta, gd t, helper t, t helper, helper, cytotoxic t, thymocyte, naive t, tcm$^w$, tem$^w$, trm$^w$, tfh$^w$, teff$^w$, th9$^w$, th22$^w$ |
| B | b cell, b-cell, b lympho, b220, cd19, cd20, igm, igd, plasma, plasmablast |
| NK | nk$^w$, natural killer |
| Myeloid | monocyte, mono, macrophage, dendritic, mono/dc, mhc, myeloid, phagocyte, apc, cd11b+ mono |
| **Fine attributes** — Capelle2023 ceiling only | |
| T coreceptor | CD8: cd8 CD4: cd4 |
| T maturation | TEMRA: temra, terminal effector, terminally differentiated; naive: naive, "tn"; CM: central memory, tcm, "cm"; EM: effector memory, tem, "em" |
| Monocyte subset | non-classical: non-classical, nc+interm, interm mono, intermediate mono; classical: classical |
| Unconventional T (fuzzy) | $\gamma\delta$: gd t, $\gamma\delta$, tcrgd, gamma delta; MAIT: mait; NKT: nkt |

Table S1: Datasets overview used in this study.

| Dataset | Samples / individuals | Disease / treatment status | Cells per sample (min–max, mean) |
|---|---|---|---|
| Capelle2023 (human whole blood) | 50 / 50 (one sample per person)[a] | 27 Parkinson's disease + 23 healthy controls | 321,545–985,782 (mean 623,591) |
| Levine2015 (human bone marrow) | 2 / 2 (donors H1, H2) | Healthy bone-marrow donors | 74,276–191,351 (mean 132,814) total; 31,721 / 72,463 manually gated |
| Samusik2016 (mouse bone marrow) | 1 / 1 (Samusik_01, C57BL/6J) | Healthy C57BL/6J mouse | 86,864 total; 53,173 manually gated |
| Parrish2023 (mouse colonic lamina propria) | 29 / 29 (mice) | Female, specific-pathogen-free BALB/c; 2 diets (fibre-rich / fibre-free) × 3 treatments (OVA-CTX, CTX, PBS); colonic lamina propria $CD45^+$ cells, 24 h post-challenge | 49,292–375,745 (mean 192,827)[b] |

[a] 50 samples remaining after quality-control exclusions from the original 52-participant discovery cohort (28 PD / 24 healthy controls) reported in Capelle et al., *Nat Commun* 2023;14:7461. [b] $CD45^+$ event counts, computed from the `$TOT` keyword of each FCS file in the Zenodo deposit (doi:10.5281/zenodo.22077384); median 173,584, 5,591,989 events in total. These are the events entering clustering. The design comprises five samples per treatment–diet arm except PBS/fibre-rich, which has four, giving 29 rather than 30 samples.

Table S2: Models evaluated, with serving configuration. Cloud models are served by the provider, so local precision and size do not apply. Local models were pulled from the Ollama library under the tags shown; precision is each tag's distribution default, except qwen3.6, which pins bf16. Parameter counts and on-disk sizes are as reported by the Ollama library.

| Model | Tag / version | Serving | Architecture | Parameters | Precision | Size |
|---|---|---|---|---|---|---|
| Claude Sonnet 4.6 | `claude-sonnet-4-6` | Cloud API | — | — | n/a | — |
| Claude Opus 5 | `claude-opus-5` | Cloud API | — | — | n/a | — |
| gpt-oss 120B | `gpt-oss:120b` | Ollama | MoE | 117B | MXFP4 | 65 GB |
| qwen3.6 35B | `qwen3.6:35b-a3b-bf16` | Ollama | MoE (3B active) | 36.0B | bf16 | 71 GB |
| gemma4 31B | `gemma4:31b` | Ollama | Dense | 31.3B | Q4_K_M | 19 GB |
| gemma3 27B | `gemma3:27b` | Ollama | Dense | 27.4B | Q4_K_M | 17 GB |
| Llama 4 Scout | `llama4:latest` | Ollama | MoE (17B active) | 109B | Q4_K_M | 67 GB |
| medgemma 27B | `symptoma/medgemma3:27b` | Ollama | Dense | 27.0B | Q4_K_M | 16 GB |

Tags, parameter counts, quantisation and on-disk sizes as reported by `ollama show` and `ollama list` on the serving host. The sizes corroborate the stated precisions: the five Q4_K_M models occupy 4.7–4.9 bits per parameter and gpt-oss 4.5, whereas qwen3.6 occupies 15.8, consistent with bf16 and with the precision mismatch noted in the Limitations.

Table S4: Unmapped-label fraction per model and dataset (%, frequency-weighted share of cells whose label matched no pattern in the dataset's keyword map; Table S3). Unmapped labels are scored as disagreements. Values track model labelling behaviour rather than map coverage: the highest belong to the frontier control Opus 5 and to the models that produce coarse or abstaining labels, and are independent of provider. Across every cell above 2%, each distinct unmapped label was a coarse term spanning several lineages, an explicit abstention, or a population absent from that dataset's reference. †Opus 5 values are the mean of the same three runs used in Tables 2 and 3 of the main text; per-run values are 0.2/0.2/0.2 (Capelle2023), 10.9/15.7/13.1 (Levine2015), 40.0/38.6/38.7 (Samusik2016) and 5.1/15.7/14.1 (Parrish2023).

| Model | Capelle2023 | Levine2015 | Samusik2016 | Parrish2023 |
|---|---|---|---|---|
| Claude Sonnet 4.6 | 0.1 | 0.4 | 0.1 | 3.9 |
| gpt-oss 120B | 0.0 | 4.9 | 4.4 | 0.2 |
| qwen3.6 35B | 0.3 | 0.5 | 2.0 | 2.7 |
| gemma4 31B | 3.3 | 19.6 | 1.9 | 14.1 |
| gemma3 27B | 0.0 | 10.8 | 1.1 | 0.0 |
| Llama 4 Scout | 0.0 | 7.4 | 22.5 | 10.6 |
| medgemma 27B | 0.0 | 1.9 | 1.4 | 0.0 |
| Claude Opus 5† | 0.2 | 13.2 | 39.1 | 11.6 |

Table S5: Metacluster-level concordance (%, frequency-weighted) under the full keyword map versus a map restricted to keywords matching a label in the dataset's own reference vocabulary (each cell: full / reference-grounded-only). The reference-only map scores standard nomenclature as unmapped whenever it differs from the reference's shorthand, so it is a synonym-blind lower bound rather than a corrected estimate. The qualitative conclusions hold under both maps: no local model leads everywhere, and Llama 4 still collapses on mouse Samusik2016. The full map does not favour Claude Sonnet. Its drop under the reference-only map is the smallest of the seven models on Parrish2023, where the restriction bites hardest (23.6 points against 33.8–63.9), and the second smallest on Levine2015, and its ranking improves from first on three datasets to first on all four. Preferential synonym coverage for that model would predict the opposite.

| Model | Capelle2023 | Levine2015 | Samusik2016 | Parrish2023 |
|---|---|---|---|---|
| Claude Sonnet 4.6 | 98.2 / 98.0 | 85.2 / 73.8 | 93.9 / 90.5 | 95.5 / 71.9 |
| gpt-oss 120B | 98.0 / 97.7 | 84.5 / 63.7 | 75.1 / 81.2 | 95.3 / 61.5 |
| qwen3.6 35B | 96.0 / 96.7 | 83.3 / 71.0 | 91.6 / 89.3 | 95.4 / 57.9 |
| gemma4 31B | 95.4 / 95.1 | 79.1 / 58.8 | 90.4 / 84.7 | 79.4 / 34.8 |
| gemma3 27B | 92.6 / 92.4 | 68.6 / 51.3 | 73.2 / 70.8 | 89.8 / 36.5 |
| Llama 4 Scout | 96.3 / 96.0 | 85.6 / 65.8 | 55.2 / 54.9 | 82.5 / 19.2 |
| medgemma 27B | 94.9 / 94.7 | 68.7 / 61.4 | 80.0 / 76.8 | 88.0 / 24.1 |
